# Innate and adaptive immune basis of mental health effect on Alzheimer’s disease progression

**DOI:** 10.1101/2022.06.09.495493

**Authors:** Yilin Feng, Jiaqi Fan, Yifan Cheng, Qionghai Dai, Shaohua Ma

## Abstract

Mental health has long been suspected to be highly associated with neurodegenerative disease, but it lacks experimental evidence to elaborate the link and immunological mechanism between them. In this work, we studied the progression of Alzheimer’s disease (AD) and its associated immune activity by using the transgenic mice under negative (depression) and positive (environmental enrichment, EE) mental health intervention. The tissue pathology, resident and peripheral immunity and behavioral characteristics were investigated. The transgenic mice undergoing depression treatment was featured with aggravated AD pathology, elevated activation of microglia in the brains, more abundance of Treg cells and cytotoxic T cells, and higher ratios of central memory T cells to effector memory T (T_CM_-to-T_EM_) cells in the peripheral blood and spleens. The EE treated mice were featured with alleviated AD pathology, reduced activation of microglia, less amounts of Treg and cytotoxic T cells and lower ratios of T_CM_-to-T_EM_ cells. This study may provide strategies for immunity regulation and mental health intervention that benefit AD therapy.

**Significance statement:** Mental health is suspected to regulate AD progression, but its validation in histology examination and the regulatory mechanism remains unclear. Here, we prove that both innate and adaptive immunity are playing important roles and mental health-immunity-AD progression forms a triad. More active microglia, Treg, and cytotoxic T cells, and higher TCM-to-TEM feature negative mental health and severer AD progress, and vice versa. The findings in adaptive immunity and the triad may inspire AD regulation.

## Introduction

Numerous studies have reported that mental health is associated with disease progression^1,2^ and causes changes in the immune network^3,4^, yet few studies have elucidated the underlying mechanism linking mental health, neurodegenerative diseases, and the immune responses. Alzheimer’s disease (AD) is one of the most common incurable neurodegenerative diseases, causing a growing problem in society for healthcare. The neuropathology of AD is mostly characterized by β-amyloid accumulation that forms plaques scattered in some brain regions^5,6^. The abundance of β-amyloid plaques grows with disease progression. Depression is a common risk of developing neurodegenerative diseases^7,8^. Major depressive disorder (MDD) is a chronic psychiatric disease that gives rise to cognitive impairment^9^, showing similar pathology to Alzheimer disease^10,11^. Patients with a history of depression may increase the risk of developing dementia^12^, though the mechanism remains unsolved. Some clinical studies reported increased hippocampal plaques in patients with a history of depression, among which, a large fraction developed apparent AD symptom^13^. High levels of glutamate, a neurotransmitter and also a marker for mood disorder^14^, were found in the cerebrospinal fluid of patients with either probable AD or depression^15^. These studies suggest the chance of depression in promoting neurodegenerative disease progression, but it remains a hypothesis and lacks experimental elucidation of the linkage and underlying cues.

On the contrary, during the last decades, environmental enrichment (EE), a behavioral intervention method to build positive emotion, has been proven to improve learning and memory^16^ in mice. EE intervention has also been reported to revert the cognitive ability of transgenic mice^17^, via a variety of effects from molecular and cellular to behavioral levels. For example, it is suggested that EE intervention elevates brain activity by causing molecular production of insulin-like growing factor (IGF-I) and brain-derived neurotrophic factor (BDNF) to augment brain function and plasticity^18^. EE accelerates the maturation of visual acuity by improving spatial localization and enhancing orientation sensitivity of primary auditory cortex (A1) neurons^18^. Other studies show that EE reduces symptoms of anxiety disorders or mental illnesses of mice, including increased exploratory activity and decreased psychological stress responses^19–22^. Though numerous studies of EE intervention have proven its positive effect on managing cognitive and mental illness, there is no conclusive study on the pathological elucidation of EE intervention on AD progression^23,24^.

Immune responses, including both the innate immune and adaptive immune activities, have been reported playing pivotal roles in AD progression^25,26^. Activation of microglia, the key innate immunity cells of central nervous system (CNS), internalizes the pathogenic species like β-amyloid and tau protein and degrades them through various endocytic pathways to modulate the neuroinflammatory process^27–29^. This process usually resolves once the immune stimulus is eliminated; however, microglia in aged brains have functional impairments and are prone to sustained activation, which might contribute to the pathogenesis of neurodegenerative diseases^30^. Abnormal activation of microglia produces neurotoxic cytokines and chemokines, and subsequently results in neuron death^29^. T cell, a major peripheral immune cell, infiltrates the brain and patrols the intrathecal space of aging or diseased brains, including brains comprising AD pathology^26,31^. Both CD8+ T cells and CD4+ T cells are involved in the AD pathology. It was reported that peripheral CD4+ and CD8+ T cells were more differentiated and pro-inflammatory in pathological tissues of AD than in normal tissues^32^. With AD progression, T cells broke through the blood-brain barrier (BBB) and entered the brain tissue to respond to antigens, leading to neuronal dysfunction and aggravation of the disease^33^.

Though observational and pathological studies have implied the triad of mental health, AD progression, and immune activity, the underlying mechanisms and dominant immunity cells remain unknown, which impedes the administration of mental health and immune modulation in AD treatment. In this study, we chose APP/PS1 mice as the AD diseased model and exposed them to depression and EE intervention as mimics of two distinct mental health conditions. It was found that depression increased microglia activation and accelerated AD progression, but EE decreased both. Notably, T cells exacerbate AD progression by increasing the abundance of total CD8+ T cells, central memory T cells and regulatory T cells (Tregs) in depression intervention, but the effect was compromised in EE mice.

## Results

### 2.1 Depression affects learning ability in AD progressed mice

Chronic social defeat stress (CSDS) has been proven to induce a depression- and anxiety-like phenotype in adolescent male C57BL/6 mice^34,35^. Here we used CSDS to establish anxiety disorder and depression models in APP/PS1 transgenic mice at the age of 6-7 weeks, following the standard protocol^35^. The CSDS modeling was executed at a young age to establish long-term depression in the transgenic mice^36–38^. The CSDS-sensitive models were screened by the social interaction test (Figure 1A & 1B). Previous CSDS study showed that the maintenance of depression-like phenotype was not life-long^39^. Thus, a 24-hour acute restraint stress^40^ was performed one month post-CSDS to prolong the depression-like behavior (Figure 1A). Two groups of mice age 6-8 months and 10-12 months were analyzed separately, because aging was also a key influence factor of AD.

**Figure 1.**
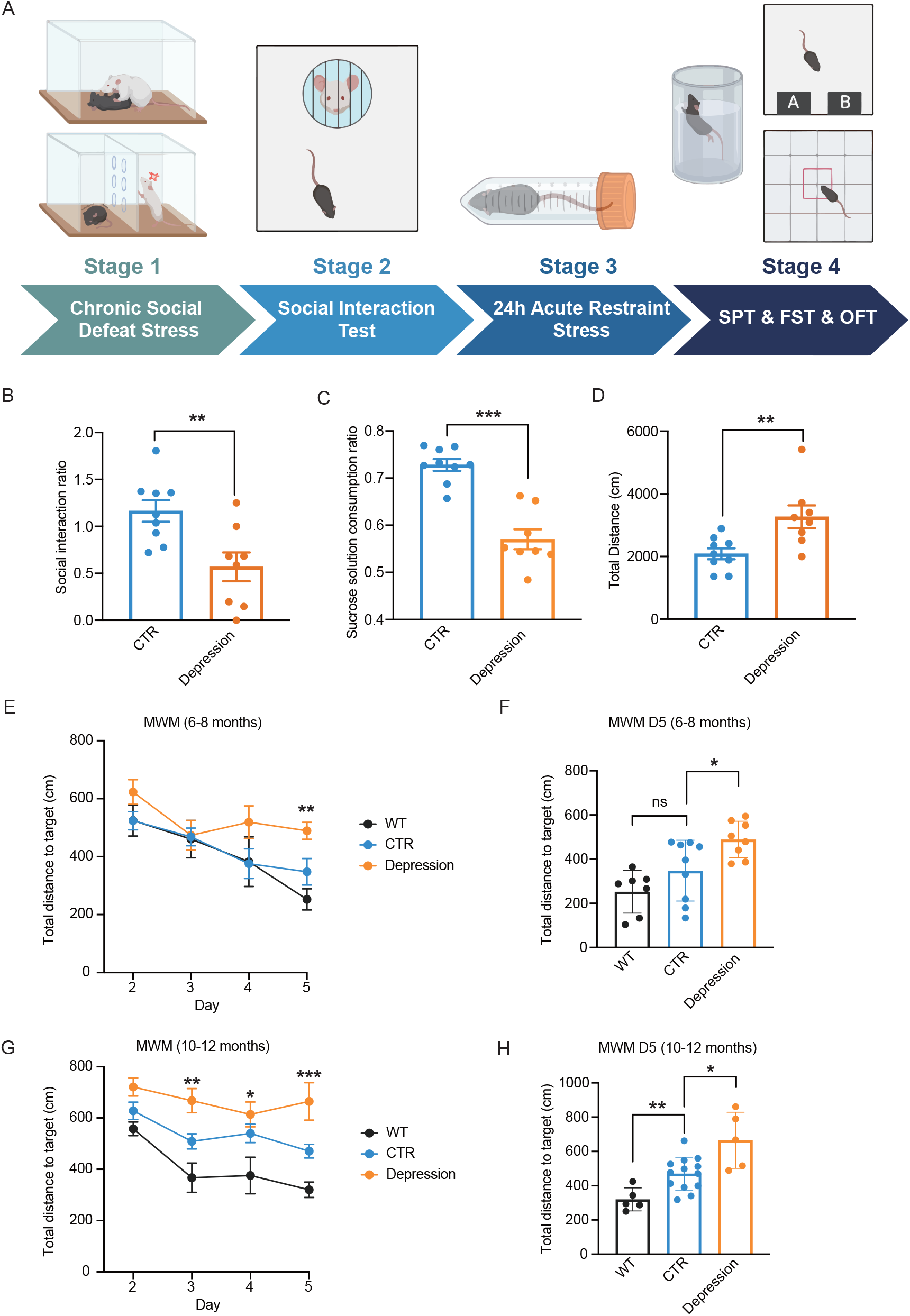
Depression modeling and behavioral test imply progressed learning deficiency in depression-modeled AD mice. A, The behavioral modeling and test process to establish and validate depression mouse models. The process contains four stages, including Stage 1-Chronic Social Defeat Stress, Stage 2-Social Interaction Test, Stage 3-24h Acute Restraint Stress and Stage 4-SPT, FST, OFT. Stage 1 builds up the social defeat depression model and is tested in Stage 2. Stage 3 reinforces the model and is tested in Stage 4. B, Social interaction ratio in Stage 2. The ratio of depression-modeled mice is significantly lower than CTR mice (p = 0.0055); C, Sucrose solution consumption ratio of SPT test in Stage 4, which shows a significant difference between depression-modeled and CTR mice. Depression-modeled mice show a lower preference for sucrose solution (p = 0.0055); D, Total distance of OFT in Stage 4, which shows a significant difference between depression-modeled and CTR mice. Depression-modeled mice show longer traveling distance in the open field, which implies obvious stress-like behavior; E, F, Total distance to the target of the mice in the MWM test. E shows the performance of the mice on days 2-5, including a learning effect and performance progress across days. Depression-modeled mice show obvious learning deficiency (p = 0.0017). F compares the performance on day 5 (p = 0.0360). B-F shows data for mice of 6-8 months of age. G and H are the same as E, F, but show the data of the mice aged 10-12 months (Left to right, G: p = 0.0015, 0.0263, 0.0003; H: p = 0.0098, 0.0264). Mann-Whitney test for two groups, Ordinary one-way ANOVA test for more than two groups; * for p < 0.05; ** for p < 0.01; *** for p < 0.001; **** for p < 0.0001;

Before proceeding to the cognitive behavioral test, we first conducted three emotional behavioral tests, namely the sucrose preference test (SPT), the forced swimming test (FST) and the open field test (OFT) (Figure 1A) to verify the depression phenotype. At the age of 6-8 months, depressive-like mice exhibited less sucrose preference (Figure 1C), and anxiety-like behavior in OFT with increased total distance (Figure 1D). There was no significant difference in FST between the depression and the control groups (Figure S1A). We used the Morris Water Maze (MWM) test, a behavioral standard for AD progression examination, to measure the learning and cognitive abilities in mice^41^. The 6-8 months old mice in the control group had no sign of cognitive decline. However, the depressed mice of the same age appeared cognitive decline, as indicated by the distance to the target platform during training from day 2 to day 5 (Figures 1E & 1F). The MWM results of the 10-12 months old mice showed that the control group had begun to loss memory, whereas the depression group, also screened by emotion behavior tests (Figure S1C – S1F), exhibited more severely impaired memory and cognitive ability (Figures 1G & 1H). It proved that depression significantly accelerated memory loss and cognitive impairment in APP/PS1 mice.

### 2.2 Environmental enrichment (EE) reverts the cognitive impairment caused by AD progression

We followed the classic method to implement EE intervention^42^. The mice were reared in enriched-environment cages for more than 2 months upon arrival at 6-7 weeks age (Figure 2A). The MWM test performed over 5 days suggested that the cognitive and learning abilities of APP/PS1 mice of the EE group were comparable to the wild-type mice even at 10-12 months old (Figures 2B & 2C). However, the control group that received no EE intervention displayed apparent cognitive decline, suggested by the increased “total distance to target” in the MWM test. Figure 2D shows the trajectories of mice movement on the test day (day 6) without the target platform. More activity near the area where the platform was placed (the target zone) on the first 5 days implied a better learning and memory ability. The EE mice and the wild-type mice moved around the target zone more than the CTR and depression mice. Figure 2E shows the numbers of times of mice crossing the target area within the tested two minutes. The EE intervened APP/PS1 mice maintained the learning and memory ability analogous to the wild-type mice, whereas both the control group and the depression group showed compromised ability. It is suggested that the learning and memory ability of APP/PS1 mice decreases with the decline of mental health, and EE intervention repairs the cognitive decline of transgenic AD mice.

**Figure 2.**
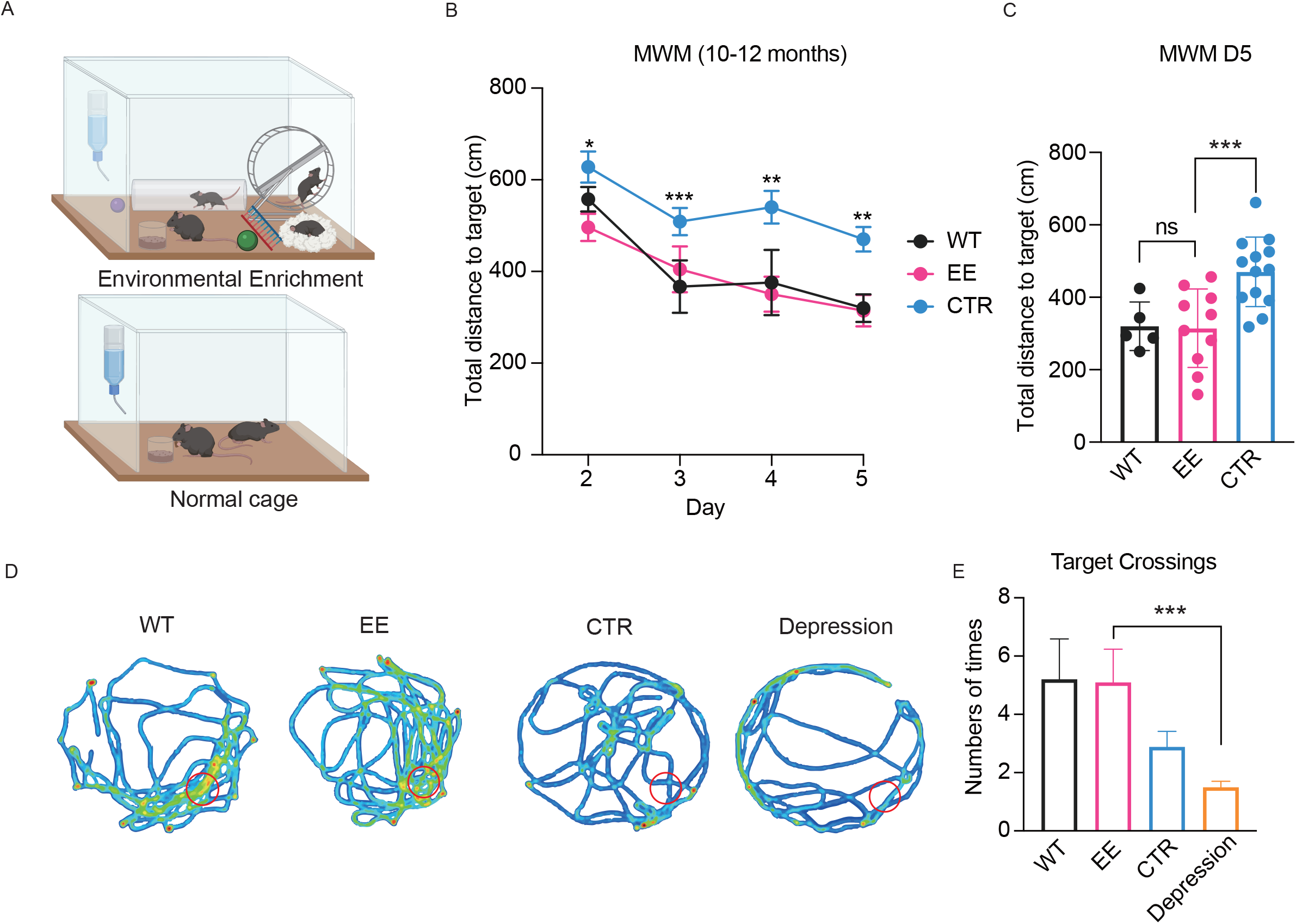
Environmental enrichment alleviates AD-caused learning deficiency. A, A schematic of environmental enrichment. B, C, Total distance to the target of the mice in the MWM test. B shows the performance of the mice on days 2-5, including a learning effect and performance progress across days. CTR mice show significant learning deficiency, while EE mice show performance comparable to WT mice (Left to right, p = 0.0208, 0.0006, 0.0049, 0.0013). C compares the performance of day 5 (p = 0.0098). D shows example traces of mice without a target in the MWM on day 6. WT and EE mice cross the target zone more than CTR and Depression mice, implying a better memory of the task. E, Statistical comparison of target-crossing of the four groups of mice on day 6 (p = 0.0007, one-way ANOVA p = 0.0337). Mann-Whitney test for two groups, Ordinary one-way ANOVA test for more than two groups; * for p < 0.05; ** for p < 0.01; *** for p < 0.001; **** for p < 0.0001;

### 2.3 Microglia-mediated innate immune in AD progression mice under different mental conditions

42- and 40-amino acid forms of β-amyloid protein are generated from the β-amyloid precursor protein (APP) by different protease activities^43^. β-amyloid 1-40 (Aβ40) and 1-42 (Aβ42) take part in the formation of amyloid fibrils in mice brains that are insoluble and resistant to degradation, and eventually result in amyloid deposits^44^. The soluble proteins in brain were extracted from 10-12 months old APP/PS1 mice to explore the expression of Aβ40 and Aβ42 in different mental health groups. The expression levels were measured by enzyme-linked immunosorbent assay (ELISA). The results showed that both Aβ40 and Aβ42 were expressed at the highest level in the depression group, whereas the mice reared in the enriched environment had the lowest expression of soluble β-amyloid protein in the brain (Figures 3A & 3B). Likewise, the highest and lowest expression of APP were found in the depression and EE group, respectively (Figure 3C).

**Figure 3.**
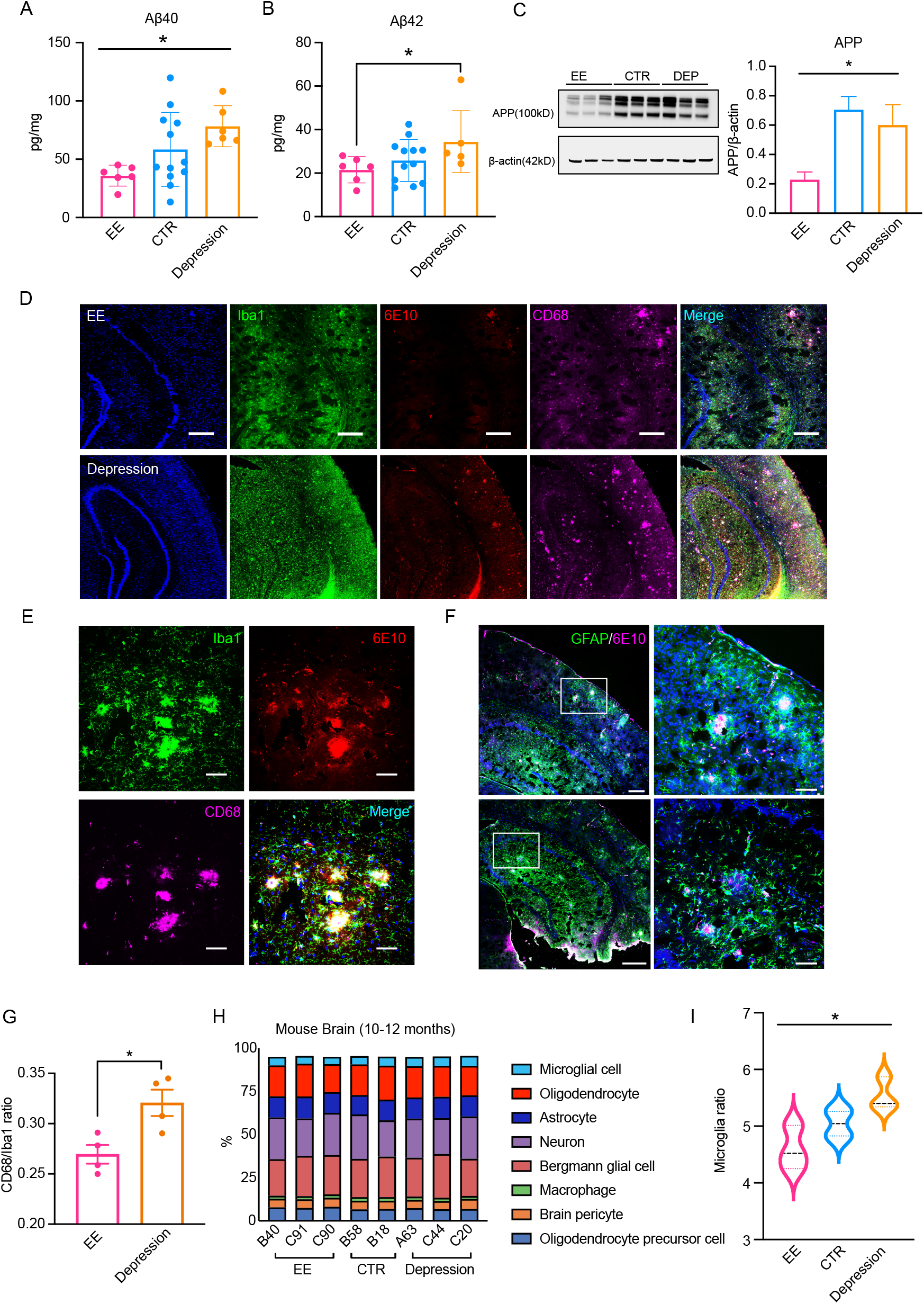
Depression and EE affect β-amyloid production and microglia activation. A, B, Elisa results of Aβ40 and Aβ42 quantification. Depression-modeled mice show more β-amyloid production, while EE mice show less (A: p = 0.0262; B: p = 0.0152); C, Western blot result of APP quantification. Depression-modeled mice show more APP production, while EE mice show less (p = 0.0345); D, Example confocal images of brain slices from EE and depression-modeled mice. The images show a part of the hippocampus, the left four images labeled by DAPI, Iba1, 6E10, CD68, and a merged image on the right. (Scale bar = 300 μm); E, Zoomed-in images of brain slices of the depression-modeled mice. (Scale bar = 50 μm); D. F, Brain slices of depression mice labeled by GFAP and 6E10, showing aggregated astrocytes around β-amyloid plaques. (First lane: scale bar = 300 μm; Second lane: scale bar = 600 μm); G, CD68/Iba1 ratio of EE and Depression mice calculated from the confocal images shows a significant difference (p = 0.0189). H, Different kinds of cells deconvolved from the bulk RNA-seq data of the brains of three groups. I, Statistical result of the microglia ratio from H (p = 0.0479). The ratio from high to low corresponds to Depression, CTR, EE, respectively. Unpaired t-test test for two groups, Ordinary one-way ANOVA test for more than two groups; * for p < 0.05; ** for p < 0.01; *** for p < 0.001; **** for p < 0.0001;

It is known that microglia cells play a role in brain immunity. Activated microglia cells internalize Aβ aggregates, but the efficiency decreases with age and pathology^45^. Conversely, microglia cells produce cytokines such as IL-1β, IL-6, TNF, and IFN-γ, which upregulate the production of β-secretase, an enzyme that cleaves APP to produce pathogenic Aβ and promote formation of amyloid plaques^29^. Therefore, aggregated microglia can lead to aggravation of AD disease. Confocal images of the brain slices from APP/PS1 mice showed that there were abundant active microglia cells, marked by Iba1, a microglia/macrophage-specific calcium-binding protein, and CD68, a surface protein of activated microglia. The activated microglia aggregated near the amyloid plaques, marked by 6E10, an anti-β-amyloid antibody (Figures 3D & 3E). Significantly higher amounts of amyloid plaques and active microglia cells were observed in the depression group (Figure 3D). In addition, astrocytes were also observed to aggregate near the amyloid foci in the mouse cortex and hippocampus (Figure 3F). Finally, the cell quantification result verified that the fraction of activated microglia was significantly higher in the depression group than in the EE group (Figure 3G).

The proportion of each type of cells in the APP/PS1 mice brains were decomposed by running the deconvolution algorithm in the bulk RNA-seq data (Figure 3H). The results showed that the proportion of microglia was significantly higher in the depression group than in the control group. The EE group gained the lowest microglia fraction (Figure 3I). The results reported that depression and EE intervention promoted or retarded the accumulation of amyloid precursor proteins and amyloid deposits, respectively, and the mental health regulation on AD progression is dependent on activation of resident microglia.

### 2.4 T cell-mediated adaptive inflammation in AD progression mice under different mental conditions

Recent studies have shown that the peripheral immune system plays an important role in AD progression, among which T cells are predominantly engaged. In AD patients^46,47^ and mouse models^48,49^, CD8+ T cells were found to be the predominant type of T cells in the brain structures related to cognition. For example, CD8+ T cells patrol the blood and cerebrospinal fluid, breakthrough BBB, infiltrate the brain parenchyma and form foci in the brains of AD patients and APP/PS1 mice^31,50^. Moreover, the ratio of CD8+/CD4+ T cells in the cerebrospinal fluid and peripheral system of AD patients is significantly higher than that of normal people^51^. Therefore, we analyzed the ratio of CD8+/CD4+ T cells using flow cytometry in the peripheral blood and the spleens of APP/PS1 mice and wild-type mice, 10-12 months old. In the peripheral blood and spleens, the highest number of CD8+ T cells, as normalized by the number of CD4+ cells (T helper cells), was recruited in the depression group than in the control, EE, and wild-type groups (Figures 4A, 4B & S2A). It suggested that the mental health condition regulated the enrichment of cytotoxic T (CD8+) cells in peripheral system.

**Figure 4.**
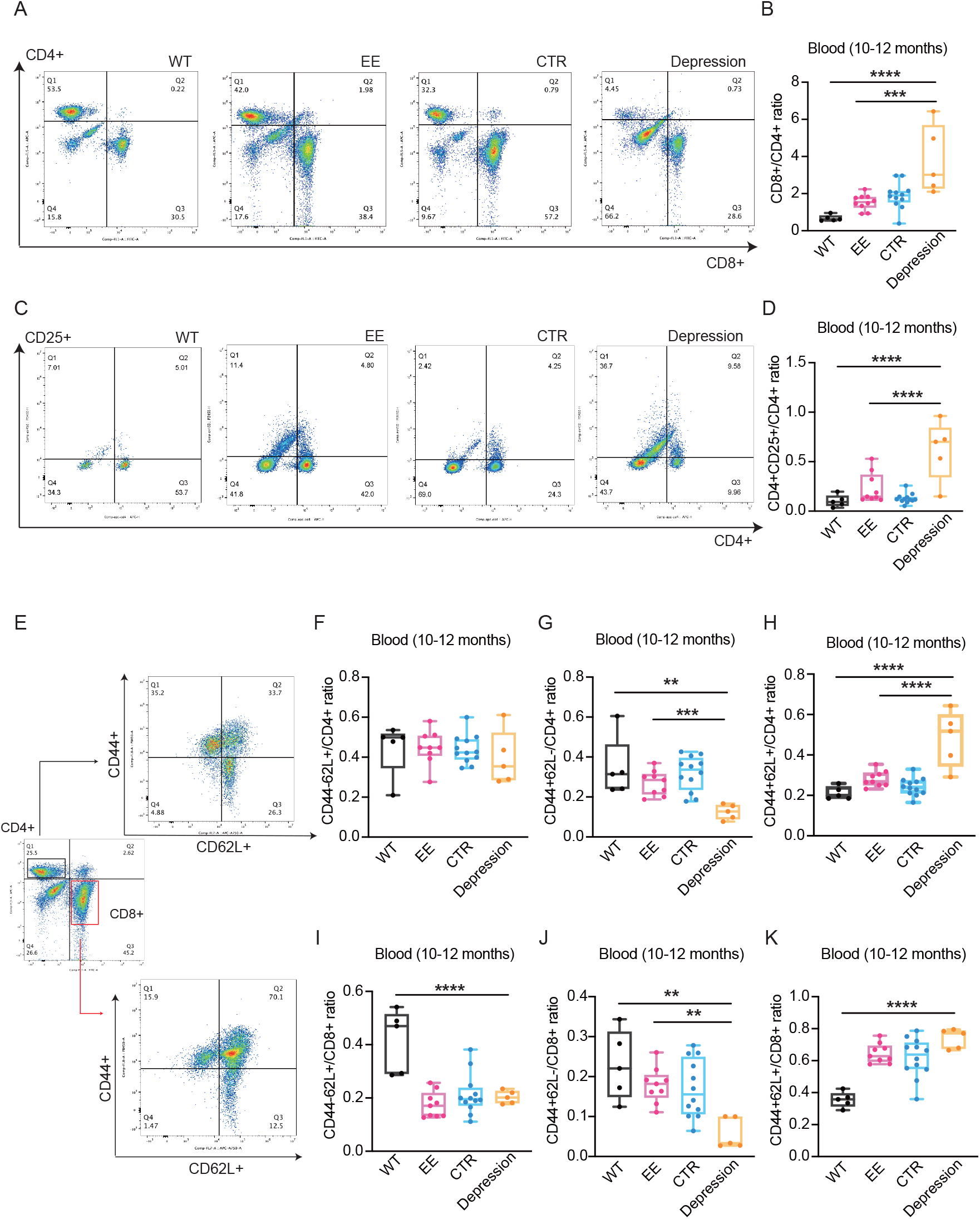
T cell populations in the peripheral blood imply different adaptive inflammation levels across groups of AD mice. A, Examples of flow cytometry plots showing the distribution of CD4+ and CD8+ T cells of one mouse from each of the four groups. B, Statistics of CD8+/CD4+ ratios in the peripheral blood of mice across four groups (4 groups: p < 0.0001; 3 groups: p = 0.0005). C, Examples of flow cytometry plots showing the distribution of CD4+ and CD25+ T cells of one mouse from each of the four groups. D, Statistics of CD4+CD25+ (regulatory T cells)/CD4+ ratios in the peripheral blood of mice across four groups (4 groups: p < 0.0001; 3 groups: p < 0.0001). E, CD44 and CD62 expression levels were quantified separately in the subgroups of CD4+ T cells and CD8+ T cells. F-H, the ratios of naive T cells (CD44-CD62L+), effector memory T cells (CD44+ CD62L-) and central memory T cells (CD44+ CD62L+) over CD4+ T cells in the blood of the mice across four groups (G: 4 groups: p = 0.0014; 3 groups: p = 0.0002; H: 4 groups: p < 0.0001; 3 groups: p < 0.0001). I-K, the same as F-H, but data from CD8+ T cells (I: 4 groups: p < 0.0001; J: 4 groups: p = 0.0018; 3 groups: p = 0.0024; K: 4 groups: p < 0.0001). Ordinary one-way ANOVA test for more than two groups; * for p < 0.05; ** for p < 0.01; *** for p < 0.001; **** for p < 0.0001;

Treg (CD4+CD25+) cells in the adaptive immune system is therapeutic for many autoimmune diseases due to their immune-regulatory functions^52^. However, in AD mouse models, the immunosuppressive activity by Treg cells may be detrimental, because it impairs the adaptive immune system that responds to Aβ pathology^53^. For example, Aβ pathology was reported significantly decreased when Tregs were depleted in either the peripheral blood or the resident tissue, whereas compounds that promoted Treg differentiation exacerbated the pathology^53^. Moreover, AD pathology was markedly elevated in immune-deficient AD-transgenic mice^54^. Therefore, the more advanced AD pathology is associated with suspected upregulation of Treg activity and vice versa. In our study, CD4+CD25+ Treg cells were identified with the highest abundance in the peripheral blood and spleens of the depression group among the four parallel groups, when normalized by the number of CD4+ cells (Figures 4C, 4D & S2B).

Immune memory is displayed by distinct T-cell subsets, including central memory T cells (T_CM_) that lack inflammatory and cytotoxic functions, and effector memory T cells (T_EM_). Mechanism studies in mice show that T_EM_ cells are more prevalent in pathological tissues, whereas T_CM_ cells are more prevalent in lymph nodes and prone to persist after infection^55^. T_CM_ cells produce IL-2 and proliferate in prompt response to antigen in secondary infection^55,56^. The ratio of T_CM_/ T_EM_ reflects the history of antigenic stimulation^56^. In inbred mice, T_EM_ transits into T_CM_ by following a linear incremental profile with time post-infection^56,57^. Alternatively, the distribution and abundance of T_CM_ and T_EM_ in pathological tissues may suggest the progression courses of AD.

Therefore, the distribution and abundance were investigated of the naive T cells (CD44-CD62L+), central memory T cells (T_CM_, CD44+ CD62L+), and effector memory T cells (T_EM_, CD44+ CD62L-) in the peripheral blood and spleens of APP/PS1 mice and wild-type mice, 10-12 months old. Subset statistics of CD4+ T cells indicated that the number of T_EM_ was significantly decreased in the depression group (Figure 4G & S2D), but the number of T_CM_ was significantly increased (Figure 4H & S2E). It suggested that the depression mice had undergone accelerated and aggravated pathological development when compared with the EE and control mice. The CD8+ T cells have analogous subset composition. Likewise, the numbers of T_EM_ and T_CM_ cells of the CD8+ T cells were the lowest and the highest, respectively, in the depression group (Figures 4J and 4K). It suggested that, in the peripheral blood and spleens, the T_EM_-to-T_CM_ conversion of CD8+ cells were the highest in the depression group, but remained lowest in the EE group (Figure S3).

Overall, a mass of cytotoxic T cell and Treg cells appeared in the peripheral blood and spleens of AD mice with negative mental health condition, which could have accelerated the progression of AD pathology. In addition, increased T_CM_ and decreased T_EM_ were observed in the T-cell immune memory pool of AD mice in the depression group, indicating that the depression group had a longer history of antigen stimulation than other parallel groups. It also proved that Aβ plaques appeared earlier in the depression group. However, the numbers of T_EM_ and T_CM_ in the peripheral blood of mice in the EE group were not significantly different from those in the control group (Figures 4D-4K). The result may suggest that delay of pathological progression of AD by EE intervention reduces recruitment of cytotoxic T cell and Treg cells.

### 2.5 Transcription profiling of depression and EE mice

Next, we performed bulk RNA-seq analysis on the brain tissues of AD mice to probe the regulatory genes on AD progression upon mental health conditioning. The depression group and the EE treatment group were compared. Figure 5C shows the top 20 differentially expressed (DE) genes between the two groups.

**Figure 5.**
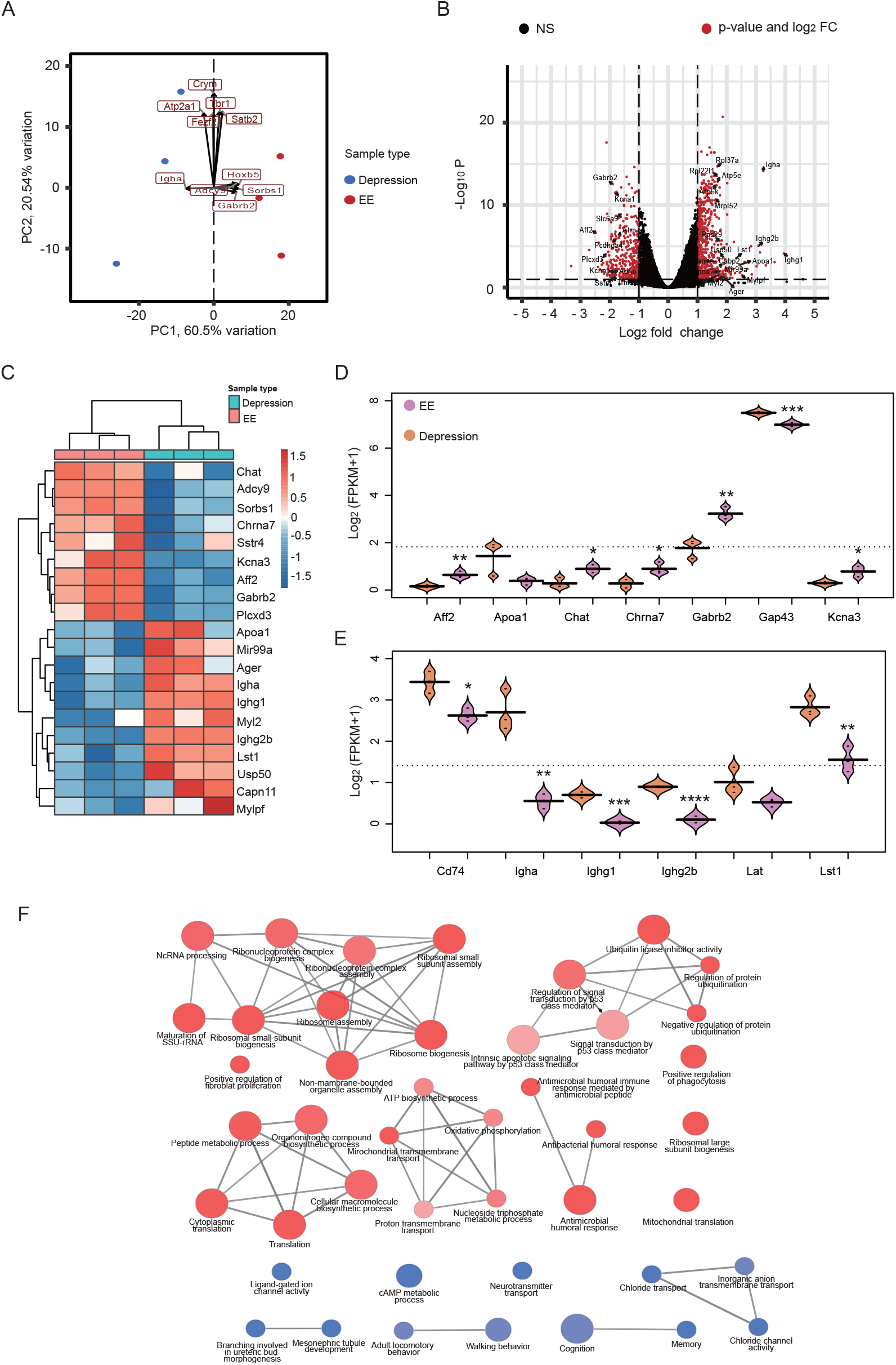
Transcriptional profiles of APP/PS1 mice under different mental health conditions. A, Principal component analysis (PCA) of gene expression for the EE group and depression group. B, Volcano plots displaying genes that are DE (adjusted P<0.01, log_2_(FC)>1) between the EE group and depression group. Red dots represent DE genes. P values were corrected for multiple testing by the Benjamini–Hochberg method. C, Heatmap of normalized gene expression in the EE group and depression group. Gene expression values were scaled by row with mean = 0 and SD = 1. D, E, Violin plots displaying significant genes between the EE group and depression group. F, Gene ontology network based on genes that are DE between the EE group and depression group. Each node represents a gene ontology. Blue nodes were based on downregulated genes in the depression group. Red nodes were based on upregulated genes.

Principal component analysis (PCA) of gene expression showed that the EE group clustered closer together and at greater distance from the depression group (Figure 5A). A total of 860 DE genes were observed between the EE group and the depression group (Figure 5B). It confirmed that transgenic mice under different mental health conditions possessed distinctive transcriptional profiles. We performed pairwise comparison between groups to screen high-rank DE genes. It was observed that the AD-related genes were highly differentiable between the two groups. Among them, cholinergic neurodegeneration was a hallmark of human AD^58,59^. The low expression of Chat (choline acetyltransferase) in the depression group was probably resulted from increased Aβ deposition that interfered the cholinergic system (Figure 5D). Both Kcna3, encoding the voltage-gated potassium channel Kv1.3, and Chrna7, encoding the α7 nicotinic acetylcholine receptor, are AD-related genes and were under-expressed in the depression group. Gabrb2 is a gene related to the GABA receptor pathway. Impaired expression of GABA signaling, i.e. under-expressed Gabrb2, in depression AD mice was observed, which might have caused E/I balance (the balance of excitation and inhibition) interruption and contributed to cognitive decline^60^. The expression of GAP-43 in AD patients has received extensive attention. The highly expressed GAP-43 in the cerebrospinal fluid is unique to AD and is associated with tau and amyloid pathology^61^. The expression of GAP-43 in the depression group was higher than that in the EE group (Figure 5D). The differential expression of afore-listed AD-related genes strengthened the correlation between mental health conditioning and AD progression.

In the former sections, we demonstrated that both innate immunity and peripheral immunity in the brain were strongly involved in the process of mental health affecting AD disease progression. Among the high-ranked DE genes, we found a large number of immune-related genes. Igha, Ighg1, and Ighg2b are immunoglobulins (Ig) related genes. The activity of the Igh family usually represents the activity of B-cell immunity^62,63^. Cd74 is involved in both T- and B-cell immune responses^64^. Both Lst, a lymphocyte-specific transcription factor, and Lat, a linker for activation of T cells, were significantly under-expressed in the EE group, suggesting the reduced activity of T cells in the brain of AD mice in the EE group (Figure 5E). In general, lymphocyte-related genes were highly expressed in the depression group but remained low in the EE group. It may suggest that the elevated adaptive immune activity appeared in AD mice of the depression group of AD mice was owing to more antigen introduction.

To investigate the function of transcriptional diversity in the two groups, we constructed a Gene Ontology network on differentially expressed genes (Figure 5F). Genes up-regulated in the depression group were associated with regulation of protein ubiquitination, ribosome assembly, oxidative metabolism, immune responses and cellular macromolecular biosynthesis processes, indicating that normal cellular processes in the depression group were disrupted, which might have promoted pathology progress. Previous studies have shown that central and peripheral immune inflammatory responses often accompany major depression disorder and metabolic syndrome^65^. Depression can lead to immune system dysfunction through glucocorticoid receptor resistance, leading to increased activation of most types of immune cells, such as microglia and effector and regulatory T cells^66,67^. The genes down-regulated in the depression group were related to cognition, neurotransmitter transmission, and chloride channel activity, indicating that the neural network homeostasis was disturbed in the depression group. Overall, it may imply the existance of the triad axis of mental health-oxidative metabolism-immunity.

### 2.6 Effect of mental condition on AD progression in female mice

The afore-presented studies were performed in male mice, including both the APP/PS1 transgenic and wild-type species. To further validate our findings and investigate whether there are latent gender differences in AD, we performed additional examination using female APP/PS1 transgenic mice. In this section, we used 21-day chronic restraint stress^68^ as the modeling method to create depression phenotype in the female APP/PS1 mice, 10-12 months old (Figure 6A). EE modeling was consistent with the method described above. The establishment of depressive-like phenotype was verified by SPT and OFT tests (Figures 6B & 6C). The MWM test showed that the depression group possessed the lowest cognitive ability, reflected by the longest time required to reach the target for the 5-day training process, whereas the EE group gained the highest cognitive ability (Figure 6D). The time spent in the target zone and the counts of target crossing were both reduced by over 3-fold from the EE group to the depression group on day 6 (Figures 6E & 6F). These findings were in accordance with the findings in the male group studies. The expression of APP protein was the highest in the depression group and the lowest in the EE group (Figure 6G). Similarly, we also analyzed the proportion of microglia cells in the brain and T cells in the peripheral blood and the spleen. The number of microglia in the depression group was higher than that in the other two groups (Figure 6H). Recruitment of CD8+ T cells and Treg cells in the peripheral blood and spleens was increased in the depression group (Figures 6I - 6J & S5A - S5B). CD8+ T_EM_ cells were reduced in the depression group while CD8+ T_CM_ cells were increased (Figures 6K – 6L), whereas the ratio of CD4+ T_CM_ and T_EM_ showed no significance among the three groups (Figures S4, S5D-S5E & S6). It was proven that the effect of mental health on female APP/PS1 mice was analogous to its effect on male mice. In addition, EE restored cognitive ability and relieved amyloid aggregation in AD progressed mice, both male and female, and improved the peripheral immunity of female diseased mice.

**Figure 6.**
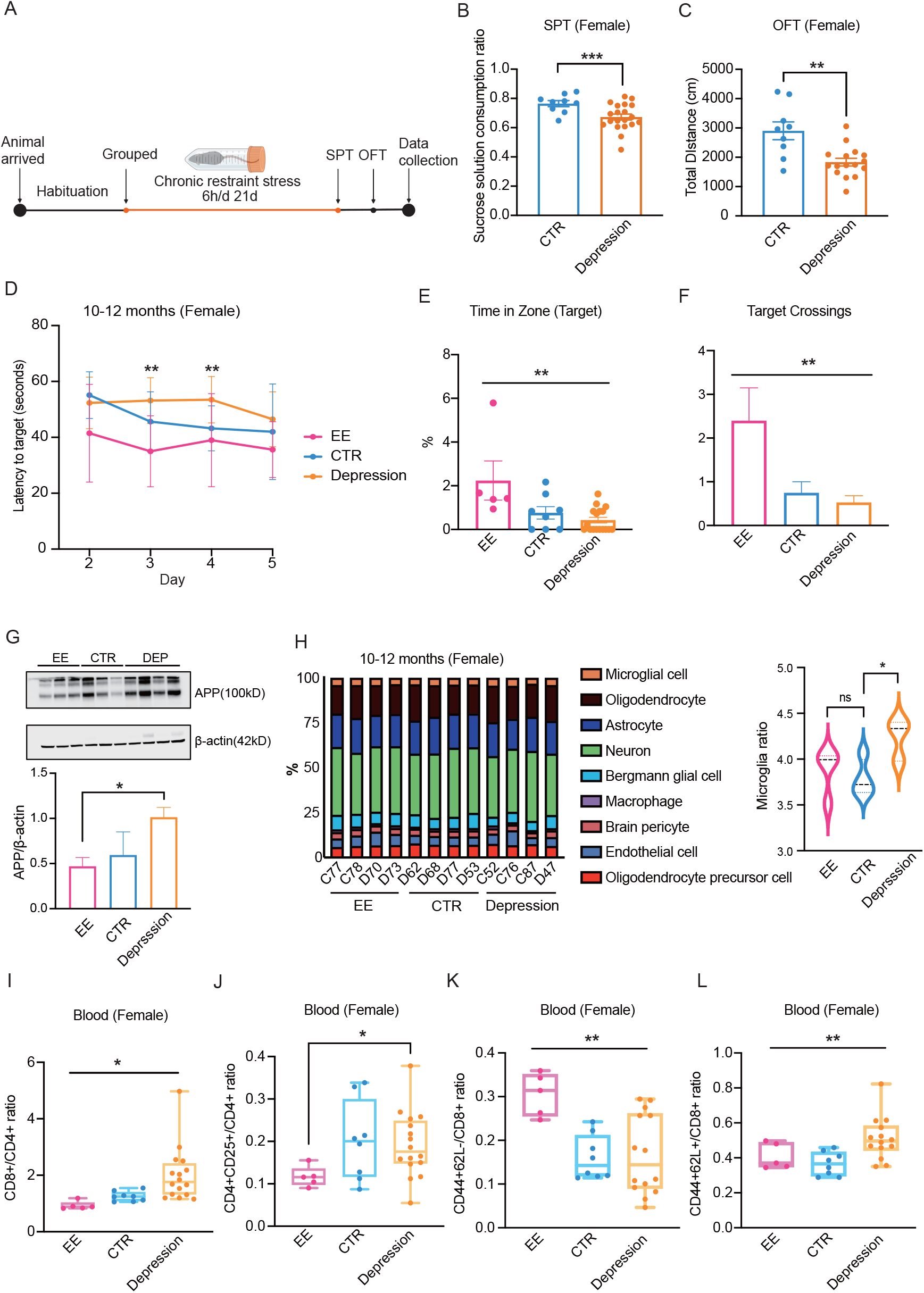
Effects of mental health condition on AD progression in female mice. A, A schematic of chronic restraint stress to induce depression in female mice. B, C, behavioral tests to validate effective depression modeling, sucrose preference test (B), and open filed test (C), respectively (B: p = 0.0060; C: p = 0.0043). D-F, MWM test showed better performance in the EE group and worse performance in the depression group. D shows latency to target on days 2-5 (Left to right, p = 0.0031; p = 0.0096). E and F show two different metrics of mice performance on day 6, time in the target zone (E) and times of target-crossing (F), respectively (E: p = 0.0040; F: p = 0.0011). G, the western-blot result shows significantly more APP expression in depression-modeled mice than EE mice (p = 0.0204). H, Left, different types of cells deconvolved from the bulk RNA-seq data of the brains of three groups. Right, a statistical result of the microglia ratio. The ratios from high to low correspond to Depression, CTR, EE, respectively (p = 0.0374). I-L, flow cytometry results of peripheral blood (I: p = 0.0102; J: p = 0.0146; K: p = 0.0037; L: p = 0.0099). Unpaired t-test test for two groups, Ordinary one-way ANOVA test for more than two groups; * for p < 0.05; ** for p < 0.01; *** for p < 0.001; **** for p < 0.0001.

## Discussion

In this study, we first explored the relationship between mental health condition and AD progression. Previous studies have shown that human beings with a history of depression have a greater risk of developing dementia. However, few studies have demonstrated whether mental health can directly affect the progression of AD disease. Here we used CSDS, 24-hour acute restraint stress and 21-days chronic restraint stress as the depression establishment methods to generate negative mental health in AD mice. In addition, we performed EE on AD mice to induce positive mental condition. The expression levels of Aβ40, Aβ42 and APP proteins in the brain and the results of MWM tests proved that depression could accelerate the progression of AD while EE could delay the onset of disease. The cognitive ability of the middle-aged EE group AD mice (10-12 months) even showed the same level as that of the wild-type mice.

We further studied the innate and adaptive immune activities in AD progressed mice under different mental health conditions to elucidate the immune basis of mental health effect on AD progression. First, depression mice featured more active microglia that accumulate alongside inflammatory lesions, whereas EE trained mice reduced distribution of active microglia, suggesting that active microglia were regulated with mental health condition and responsible for AD progression. It is in line with the senario that aggregated active microglia tend to cause neuronal death and amyloid deposition in the brains of AD patients^29^. In adaptive immunity, we observed that the ratio of CD8+/CD4+ in peripheral blood was much higher in the depression mice than in the other two groups, suggesting that depression would recruit more cytotoxic T cells in the peripheral system. In addition, Treg cells were also highly expressed in the depression group. The higher enrichment of Treg cells was suspected of compromising the gateway for leukocyte trafficking to CNS and ultimately promoting the pathology progress of AD mice^69,70^. The T_CM_-to-T_EM_ ratios of CD4+ and CD8+ cells in different groups suggested that the T cell immune memory responded differentially to mental health conditions. It was found that, upon earlier exposure to Aβ antigen, more T_CM_ but not T_EM_ appeared in the depression group with aggravated disease progress. Here, we attributed the accelerated progression of AD in depression mice to the elevated active microglia in brain, total CD8+ T cells, Treg cells and higher ratio of T_EM_-to-T_CM_ cells in the peripheral system. However, the dynamic changes of T_CM_ and T_EM_ in mice were rather complicated, and we didn’t provide the origin of memory T cells. Clarifying the mechanism may provide a better understanding of the changes of adaptive immunity in AD progression caused by mental health conditioning.

Adaptive T cell therapy is now gaining incremental attention in neurodegenerative diseases. Though it was reported that Treg cells impeded progression of AD^32,71^, other studies reported different findings^69,72^. Our study supports the positive correlation of AD progression and Treg abundance, and implies that the two factors adopt a mutual-promotion relationship. In addition, judging from the inverse correlation of T_EM_ abundance and AD progression, it is prospective to target T_EM_ cells in AD therapy or sustainable management. For this purpose, it demands further exploration to apprehend T_EM_ and Tregs functions via mental health conditioning. We suggest that positive mental condition could boost systemic immunity in neurodegenerative disease.

Here we propose a triangular relationship diagram (Figure 7). Mental health disorders, e.g. depression, can accelerate disease progression in AD mice, including increased expression of APP and Aβ protein, decreased cognitive ability, and increased plaque accumulation in the brain tissues. In addition, depression changes innate and peripheral immunity by causing changes in systemic oxidative metabolic pathways (Figure 5F). In turn, the altered immunity further accelerates neuron degeneration resulting from cytotoxic T cells and aggregation of active microglia and amyloid plaques. Further, we propose that mental health disorder not only triggers the activation of microglia and CD8+ T cell^66^, but also triggers the occurrence of Tregs^67^, which causes dysfunction of effector immune cells^73^ and paves the way for Aβ accumulation. Consequently, pathology progress exacerbates the population of CD8+ T cells, resulting in both early-onset and accelerated progression of AD characteristics. Positive mental intervention plays opposite roles in regulating the immune responses and pathology development.

**Figure 7.**
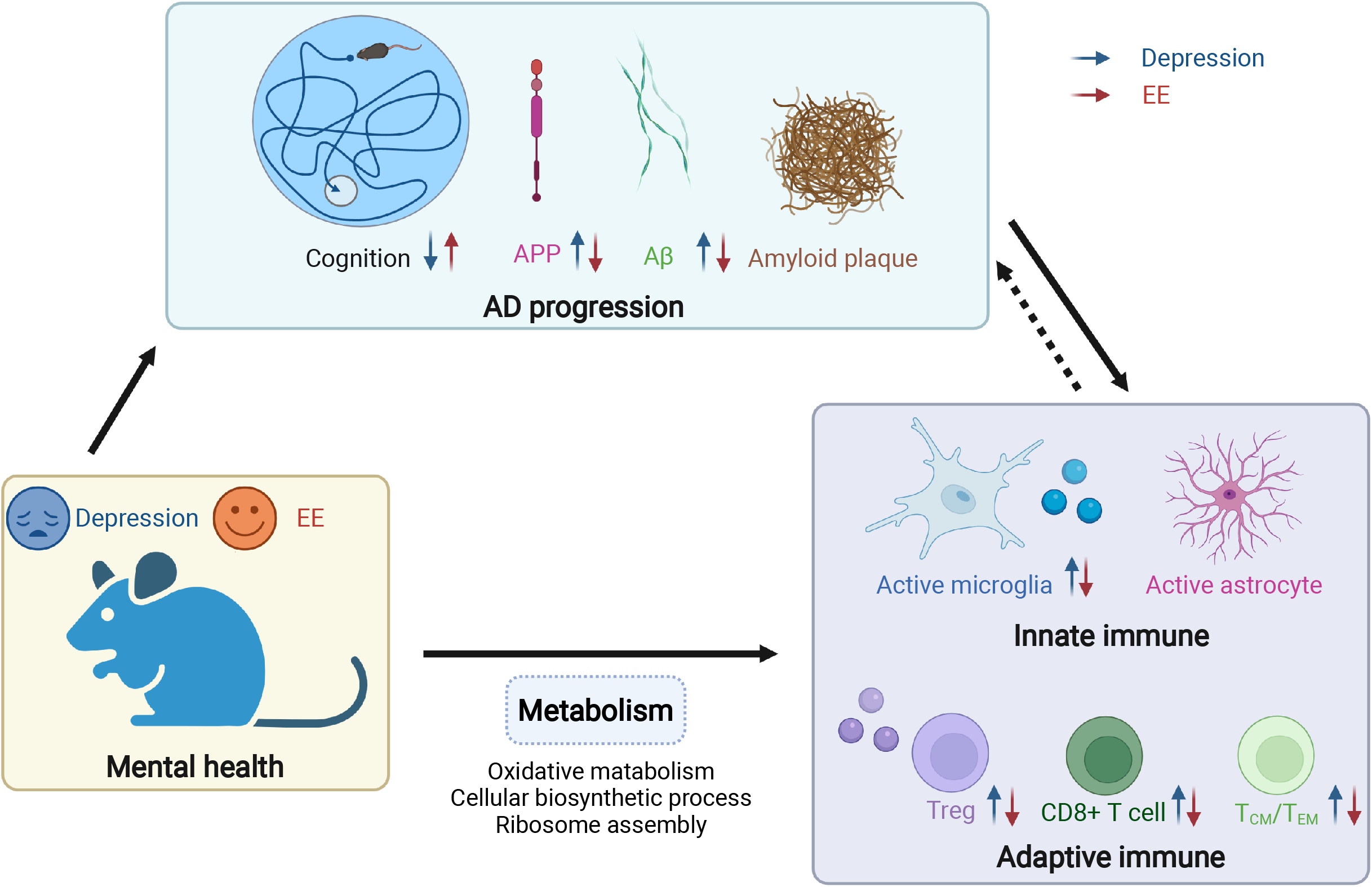
The triad linking mental health, AD pathology and immunity. The red and blue arrows represent the changes of various factors in AD mice during EE and depression, respectively. Depression results in accelerated disease progression, including cognitive decline, the increased expression level of Aβ and APP proteins, and the accumulation of amyloid plaques in the brain. Depression also changes innate and adaptive immunity by affecting the metabolism pathway in AD mice. Adaptive immune changes include increased CD8+ T cells, decreased Tregs, and increased T_CM_/T_EM_ ratios. Innate immune changes in the brain include active microglia and astrocytes. In addition, AD progression and changes in the immune network have mutual interaction. EE performs opposite functions in AD mice.

## Methods

### Animals

4 months old female and male C57BL/6 and APP Swedish PS1 dE9 mice expressing a chimeric mice/human mutant amyloid precursor protein (Mo/HuAPP695swe) and a mutant human presenilin 1 (PS1-dE9) both directed to CNS neurons under the prion protein promoter, were bought from BEIJING HFK BIOSCIENCE. 6 months old CD-1 mice were bought from Charles River. Mice were reared under standard conditions at a temperature of 22 °C and a 12 h light/dark cycle with access to standard food and water. All the animal procedures were approved by the Animal Experimentation Ethics Committee at Tsinghua Shenzhen International Graduate School.

### Depression model establishment

We followed the classic CSDS model^35^. Briefly, CD1 male mice with consistent challenge latency (≤30 s for three consecutive screening tests) were housed individually in cages fitted with perforated plexiglass partitions. The barrier blocked physical contact but allowed sensory contact. Mice aged 1-2 months were exposed to such social defeat stress for up to ten minutes per day and housed in a compartment adjacent to the attacker for the rest of the day. This process lasted for ten days, and each experimental mice would not encounter a second attack by the same attacking mice during these ten days.

According to the 24-hour acute restraint model established by Chu et al.^40^, each transgenic mice were restrained in a 50ml tube for up to 24 hours. During this period, 50ml tubes were placed in a dark box, shielded from outside light and sound. Experimental mice endured a day without food and water, which mimics the depression in humans after a disaster.

As for depression establishment on female mice, here we used 21-day chronic social defeat stress^68^. In short, female mice were restrained in 50 ml tubes in the dark for up to six hours (14:00-20:00) each day and housed in their original standard cages for the remainder of the day. This step was repeated for 21 days.

### Social interaction test

The social interaction test was used to assess social avoidance in mice following the CSDS model. Before the test, experimental mice were placed in an open area to habituate the environment for an hour. The mice were subsequently removed. Place an empty wire cage in the center of the area. In the first 150 seconds, the experimental mouse explored the area of the wire cage without the target. The distance and time of the mouse spent in the social interaction area were recorded. Then removed the mice and placed it aside. A screened novel CD1 mouse was placed in the wire cage within 30 s. During the second 150 seconds, the experimental mouse was repositioned back into the area, with the wire cage in the center of the area containing CD1 mouse. All data was collected in an automated manner by the video-tracking apparatus and software for all test phases.

### Sucrose preference test

Anhedonia is one of the core symptoms of depression. It is known that the sucrose preference test can be used to measure anhedonia in mice, following traditional protocols. Briefly, in the first 48 hours, mice were acclimated to two bottles, one with water and the other with 1% sucrose solution. During the acclimation period, the positions of the two bottles need to be changed once to prevent the mice from developing positional preference. Baseline measurements were taken after acclimatization. Mice were deprived of water and food for 24 h. After deprivation, two water bottles (water/sucrose solution) and food were given. The weights of two bottles were recorded the next day (24 hours later). During the 24 hours of the sucrose preference test, the positions of the two bottles needed to be changed every 12 hours.

### Open field test

Open field test was used to assess locomotor and anxiety-like behavior in mice. The open-air maze consists of a walled area with sufficient height to prevent subjects from escaping. Bring the mice in their home cages from their housing room to the testing room. Allow the mice to acclimate to the operating room for at least 30 min before starting the test. Gently grab its tail, remove one mouse from its cage, and place the mouse in the middle of the open maze. Allow the subject mouse to move freely and uninterruptedly within each quadrant of the maze for 10 min, during which time the tracking software will record the movement. At the end of the testing period, gently pick up the subject mouse, remove it from the maze and return it to its home cage.

### Forced swim test

The forced swim test (FST), which is one of the most commonly used tests to study depressive-like behavior in rodents. FST is based on the assumption that when an animal is placed in a container full of water, it will first try to escape, but will eventually show immobility, which may be thought to reflect a degree of behavioral desperation. The transparent cylindrical glass container has a height of 50 cm and a diameter of 20 cm. Fill the cylinder with 25 °C tap water and adjust the water depth according to the size of the mouse so that its hind legs cannot touch the bottom of the container. For mice - have a 6 min long session divided into pre-test (first 2 min) and test (last 4 min). Turn on the camera and place each mouse in a water-filled cylinder container for 6 min. Change the water after each session to avoid any impact on the next mouse.

### Environmental Enrichment

Referring to traditional EE, upon arrival of 6-8 weeks old APP/PS1 mice, control groups were housed in standard cages, each with 5 residents. Whereas, mice reared in the enriched environment were housed in a large cage of size (475*350*200mm) consisting of two running wheels, two shelters, some molar branches and tunnels. One EE cage houses 10-15 mice. The placement of the facilities will be changed every two weeks, in order to give the mice sensory stimulation.

### Morris Water Maze

Morris Water Maze (MWM) is widely used to study spatial memory and learning. The animals were placed in an opaque pool colored with skim milk powder, where they had to swim to a hidden escape platform. For the water maze scheme, we referred to the protocol of Vorhees et al.^41^. Specifically, on the first day, mouse was placed in the starting position and the time required to find the platform was recorded. The platform is marked and exposed upon the water. If the mouse did not find the platform within 1 min, it was guided to the marked platform and stayed for 15 s. The initial position is set to five. The interval between each trial was 30s. From the second to the fifth day, the water level was raised to cover the platform. Mice still started from five different initial positions, and the rest of the process was the same as the first day. During four days of the training period, the order of the five initial positions varied. On the sixth day of the test, the platform was removed, mouse was placed at the starting position farthest from the platform, and the distance and time of the mouse’s movement in different quadrants in the water maze were recorded within 2 min.

### Perfusion and surgery

After behavioral testing and peripheral blood flow cytometry, mice were gas anesthetized with isoflurane. Anesthetized mouse was perfused systemically with 20 mL of ice-cold PBS transapically to ensure high cell viability. The abdominal cavity was opened and the splenectomy was performed for flow cytometry of splenocytes. Then the mouse brain was immediately removed. The posterior half of the left hemisphere was isolated and fixed in ice-cold 4% paraformaldehyde for overnight for immunofluorescence staining. The posterior half of the right hemisphere is used for lysing by RIPA lysate to extract brain proteins. The brain stem and the anterior half of the brain were used to extract RNA for RNA sequencing.

### Flow cytometry analysis

Fresh anticoagulated whole blood was taken from the mouse eye canthus with an anticoagulant tube, and the whole blood was diluted with an equal volume of PBS. The buffy coat layer (lymphocyte layer) was separated by centrifugation at 500 g for 25 minutes at room temperature using Ficoll separation solution. The buffy coat cells were gently pipetted in a 15ml centrifuge tube and washed with 10mL PBS (1×). The isolated lymphocytes were incubated with the antibody (PE anti-mouse CD3 Antibody, PE Rat IgG2b, κ Isotype Ctrl Antibody, Pacific Blue™ anti-mouse/human CD44 Antibody, Pacific Blue™ Rat IgG2b, κ Isotype Ctrl Antibody, APC/Cyanine7 anti-mouse CD62L Antibody, APC/Cyanine7 Rat IgG2a, κ Isotype Ctrl Antibody, from Biolegend, US; EV450 Anti-Mouse CD25 Antibody [PC-61.5.3], EV450 Rat IgG1, κ Isotype Control [HRPN], from Elabscience, China) for 40 minutes on ice. The cells were filtered with a 300-mesh Falcon centrifuge tube before loading. All flow cytometry data were analyzed using Flowjo.

### Immunofluorescence

Mouse hemibrains were fixed overnight in 4% paraformaldehyde at 4°C and dehydrated in 30% sucrose solution in PBS. Tissue was embedded in OCT and frozen at −20 degrees to form a block. After making frozen blocks, put them into a constant temperature cryostat for continuous frozen section (20um). Sections were air-dried at room temperature for one hour and washed three times with PBS for 10 min each. After blocking with blocking solution (5% donkey serum, 1 × PBS, 1% triton) for one hour, the sections were incubated with primary antibodies (anti-6E10 (Biolegend, US); anti-Iba1 (WAKO, JP); anti-CD68 (Invitrogen, US); anti-GFAP (Sigma, US) diluted in blocking solution overnight at 4°C. Wash three times with PBST (1 × PBS, 1% triton) the next day. Sections were incubated with secondary antibodies for three hours at room temperature, and after three washes, nuclei were counterstained with DAPI. Sections were observed under a confocal microscope (Nikon).

### Bulk RNA-seq analysis

In order to remove technical sequences, including adapters, polymerase chain reaction (PCR) primers, or fragments thereof, and quality of bases lower than 20, pass filter data of Fastq format were processed by Cutadapt (v1.9.1)^74^ to be high quality clean data. Then clean data were aligned to the reference genome (mm10) using the software Hisat2 (v2.0.1)^75^ and counted by HTSeq (v0.6.1)^76^. Samples were subsequently analyzed using R/Bioconductor. The Rpackage DESeq2^77^ was used to normalize the data and to perform differential expression analysis. For analysis of differential expression genes, we applied a stringency level where the adjusted P value was equal to 0.05 and log_2_FC was less than −1 or >1. FC is the fold change.

### ELISA & Western blotting

Brain tissue was homogenized in RIPA (Radioimmunoprecipitation assay) lysis buffer, and the homogenate was transferred to ice for ten minutes, during which time it was shaken vigorously 3-4 times for 30 sec each time. Centrifuge at 12000 rpm for 5 min at 4 degrees, and take the supernatant for the subsequent western blotting and ELISA operations. The supernatants were collected and quantified for soluble Aβ40 and Aβ42 using enzyme-linked immunosorbent assay (ELISA) kits (Invitrogen, US) according to the manufacturer’s instructions.

The expression levels of APP and β-actin were analyzed by Western blotting. Heat the protein samples with 4× loading buffer and β-mercaptoethanol at 75 degrees for 10 min. Protein samples were separated by 4% - 20% gradient precast gels and transferred to PVDF (polyvinylidene difluoride) membranes. Membranes were incubated with 5% nonfat dry milk in Tris-buffered saline containing Tween (TBST) (10 mmol/L Tris, pH 7.5, 150 mmol/L NaCl, 0.05% Tween-20) and treated with the corresponding primary antibody (anti-β-actin and anti-APP; Cell Signaling Technology, Boston, USA) overnight at 4 °C. Protein bands were quantified by densitometry (Biorad, USA) after incubation with horseradish peroxidase-conjugated secondary antibodies for 1 hour at room temperature.

### Transcription profile analysis

To predict putative biological functions based on differentially expressed genes, we carried out a gene ontology enrichment analysis. Genes that were differentially expressed (adjusted P<0.05) between the EE group and the depression group were used to perform GO enrichment analysis. All genes were inserted into ClueGO^78^ (v2.5.8) to obtain enriched ontologies (P<0.05).

### Assessment of cell-type proportions

Scaden^79^ was used for cell-type deconvolution of our bulk RNA-seq dataset. As reference samples, we used Tabula Muris^80^, a single cell transcriptome dataset. Next, we performed scaden to calculate the cell-type proportions.

### Confocal image analysis

The brain slices were characterized by immunofluorescence. Microglia were labeled by the anti-Iba1 antibody conjugated with FITC. Activated microglia were labeled by the anti-CD68 antibody with PerCP-Cyanine5.5. The confocal images of the brain slices were analyzed with MATLAB R2021a. First, a Gaussian filter was used to filter out some noise and smooth the images. The radius of the Gaussian filter was 30 for the FITC channel, and 100 for the PerCP-Cyanine5.5 channel. The parameter sigma was set to 50 to acquire high-quality images. After that, the images were binarized with a uniform threshold. The numbers of pixels with non-zero values in the binary images were computed and the ratio of the numbers between the two channels were obtained. The ratio was used as a metric of the activity of microglia.

### Statistical analysis

All statistical analyses were performed using commercially available software (Prism, SPSS or Excel). All values are expressed as mean ± SEM. For analysis of difference between multiple groups, use one-way analysis of variance (ANOVA) to analyze the difference in means across multiple data sets. For the comparison between the two groups, Mann-Whitney test and unpaired t-test were used for significance analysis. No statistical methods were used to predetermine sample size. All graphs were drawn by using AI and biorender.

## Acknowledgements

The work was supported by the National Natural Science Foundation of China (Grant Number: 61971255 and 82111530212), the Natural Science Foundation of Guangdong Province (Grant Number: 2021B1515020092), the Shenzhen Science and Technology Innovation Commission (Grant Number: WDZC20200821141349001, RCYX20200714114736146, KCXFZ20201221173207022, KCXFZ20200201101050887), and the Shenzhen Bay Laboratory Fund (Grant Number: SZBL2020090501014).

## Author Contributions

Y.F. and J.F. contributed equally to this study. Y.F. and J.F. performed the experimental work and data analysis. Y.C. contributed to the bulk RNA-seq analysis. S.M. Q.D. Y.F. J.F. and Y.C. contributed to paper writing and revising. S.M. and Q.D. conceived and advised on this work. All authors contributed to discussion.

## Conflict of Interest

The authors declare no competing interests.

---

**Figure S1.**
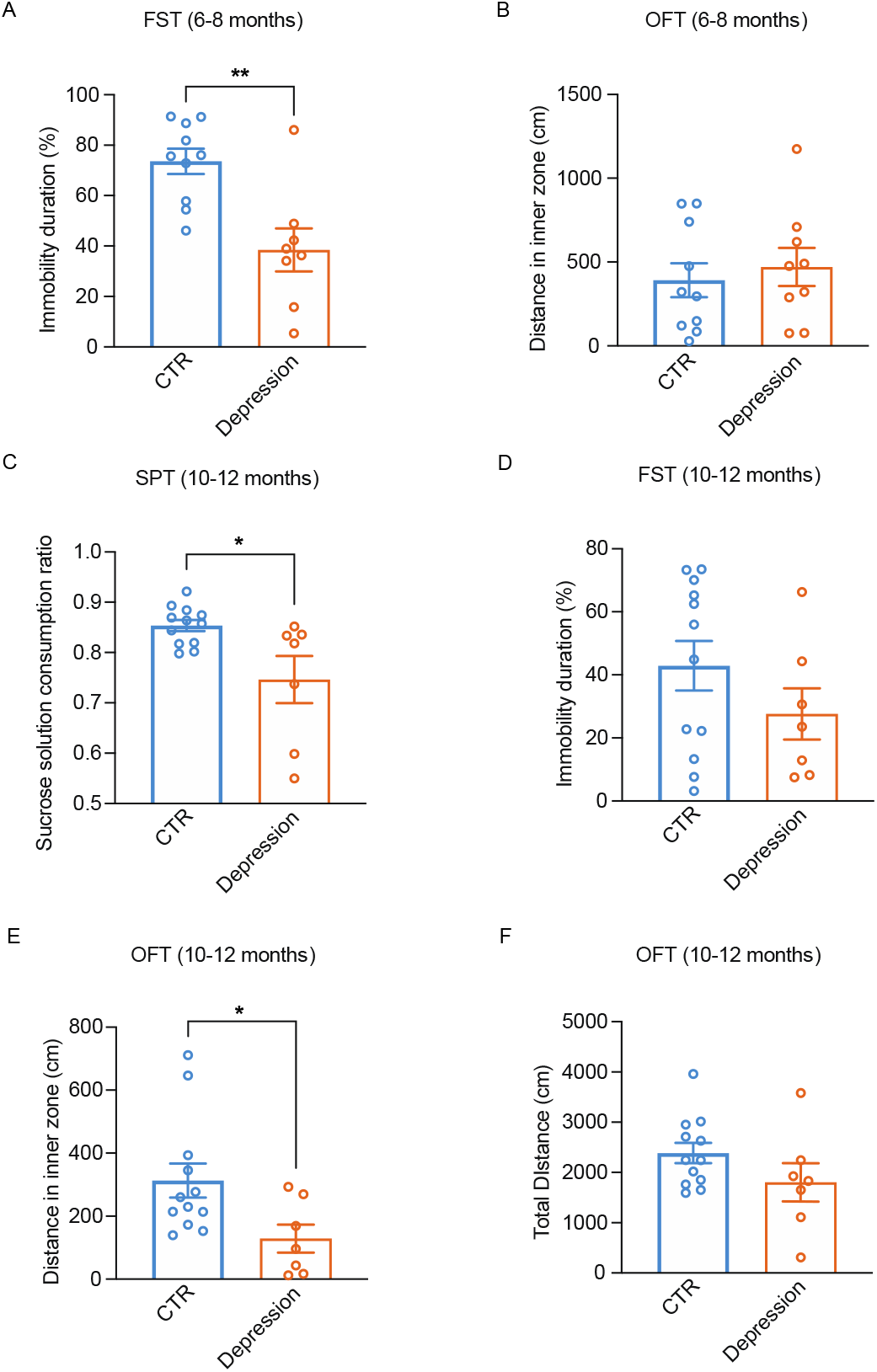
Emotion behavior tests. **A,** FST for 6-8 months mice. There is no significant difference in immobility time ratio between the control group and the depression group. **B,** OFT for 6-8 months mice. There is no significant difference in the movement distance in the inner zone between the two groups. **C,** Sucrose solution consumption ratio of SPT test, which shows significant difference between depression-modelled and CTR mice (10-12 months). Depression-modelled mice show lower preference to sucrose solution. **D,** FST for 10 - 12 months mice. There is no significant difference in immobility time ratio between control and depression groups. **E, F,** OFT for 10-12 months mice. **E,** Inner zone distance of mice OFT, which shows significant difference between depression-modelled and CTR. Depression-modelled mice show longer travelling distance in the inner zone **F,** Total distance of OFT, which shows significant difference between depression-modelled and CTR mice. Depression-modelled mice show longer travelling distance in the open field, which implies obvious stresslike behavior. Mann-Whitney test for two groups, * for p < 0.05; ** for p < 0.01; *** for p < 0.001; **** for p < 0.0001.

**Figure S2.**
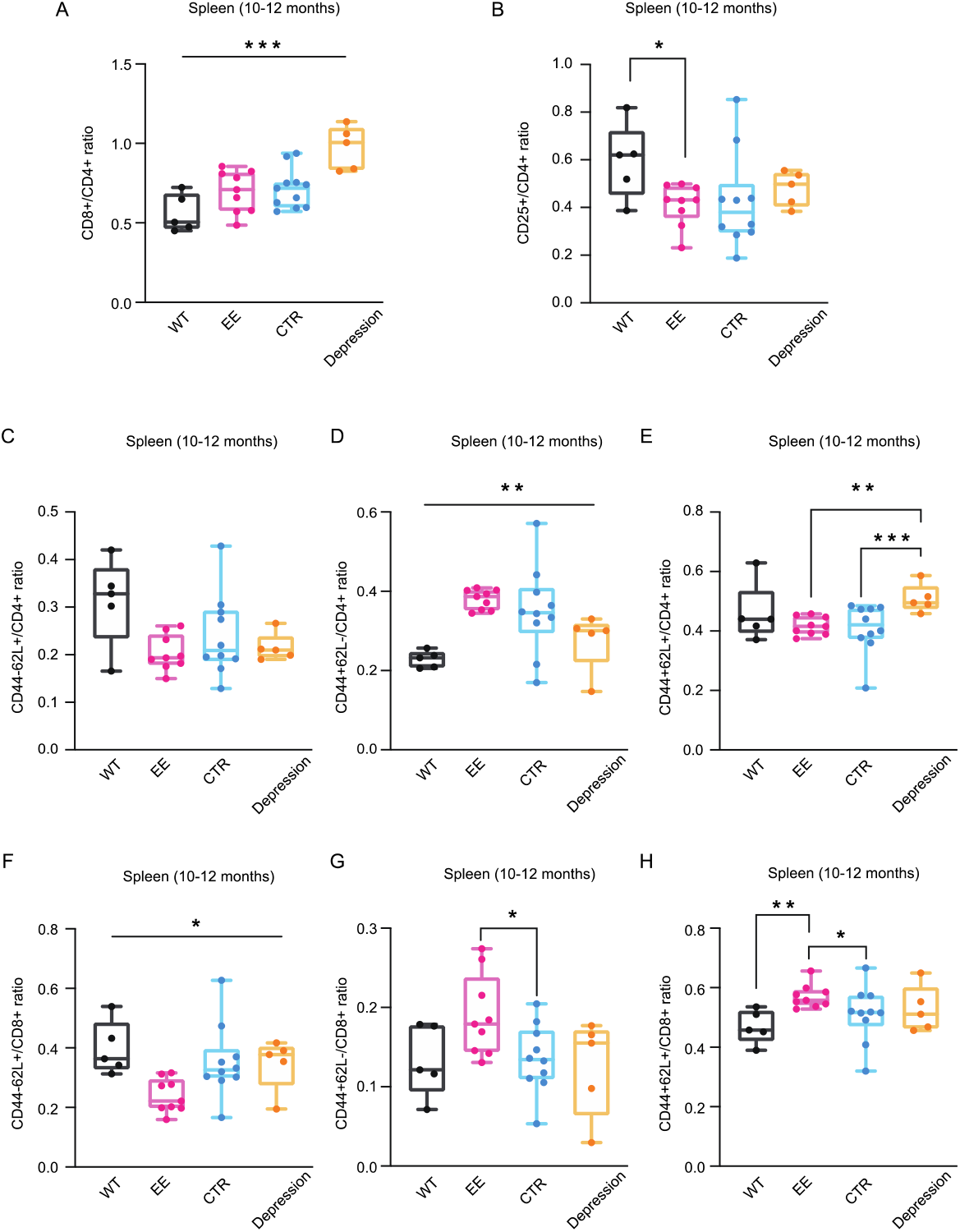
T cell populations in spleens of 10-12-month male mice. **A,** Statistical results of CD8+/CD4+ ratio in the spleen of mice across four groups. **B,** Statistical results of CD4+CD25+ (regulatory T cells)/CD4+ ratio in the spleens of mice across four groups. **C-E,** the ratio of naive T cells (CD44-CD62L+), effector memory T cells (CD44+ CD62L-) and central memory T cells (CD44+ CD62L+) of CD4+ T cells in the spleens of the mice across four groups. **F-H,** the same as **C-E,** but the data was quantified from CD8+ T cells. Ordinary one-way ANOVA test for more than two groups; * for p < 0.05; ** for p < 0.01; *** for p < 0.001; **** for p < 0.0001.

**Figure S3.**
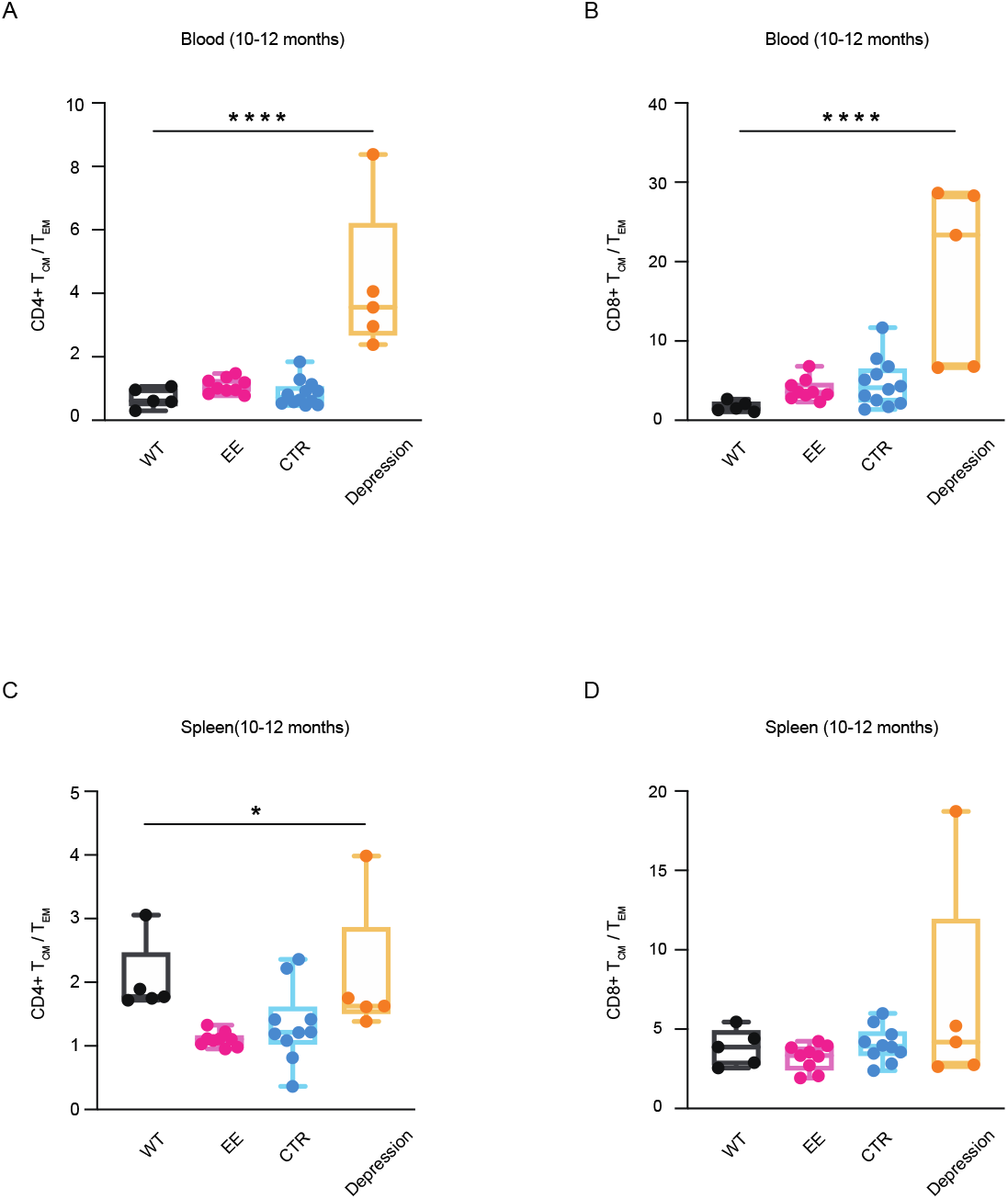
T_CM_-to-T_EM_ ratio of 10-12-month male mice. **A,** CD4+ T_CM_-to-T_EM_ ratio of peripheral blood. **B,** CD8+ T_CM_-to-T_EM_ ratio of peripheral blood. **C, D,** the same as **A, B** but statistics for spleens. Ordinary one-way ANOVA test for more than two groups; * for p < 0.05; ** for p < 0.01; *** for p < 0.001; **** for p < 0.0001.

**Figure S4.**
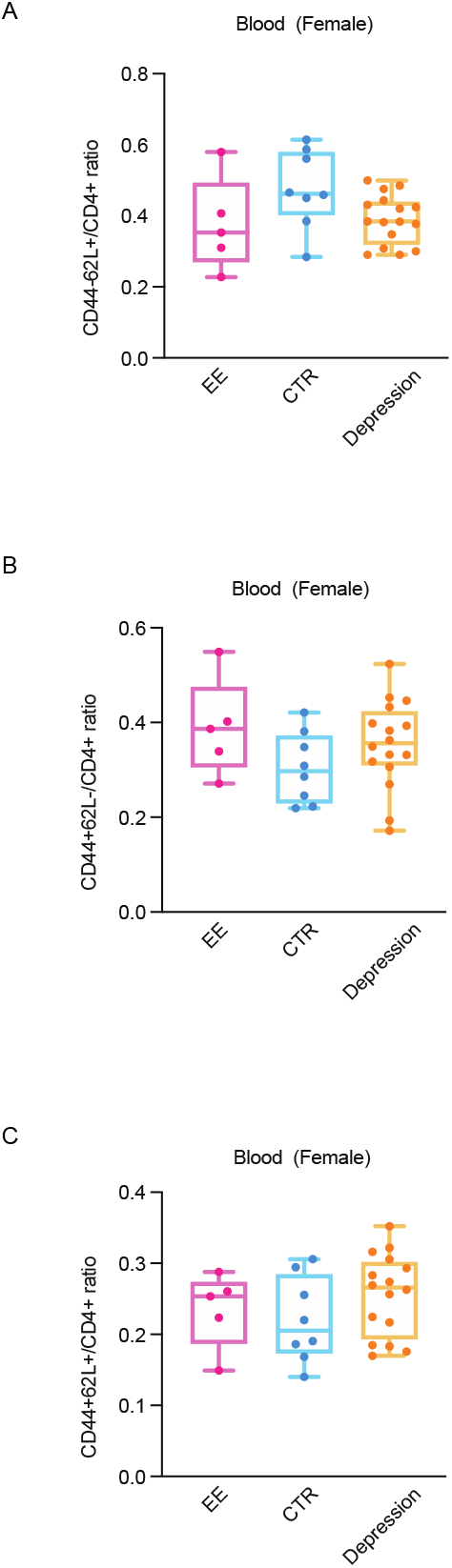
CD4+ memory T cell populations of peripheral blood in 10-12-month female mice. **A-C,** The ratio of naive T cells (CD44-CD62L+), effector memory T cells (CD44+ CD62L-) and central memory T cells (CD44+ CD62L+) of CD4+ T cells in the peripheral blood of the female mice across three groups. Ordinary one-way ANOVA test for more than two groups; * for p < 0.05; ** for p < 0.01; *** for p < 0.001; **** for p < 0.0001.

**Figure S5.**
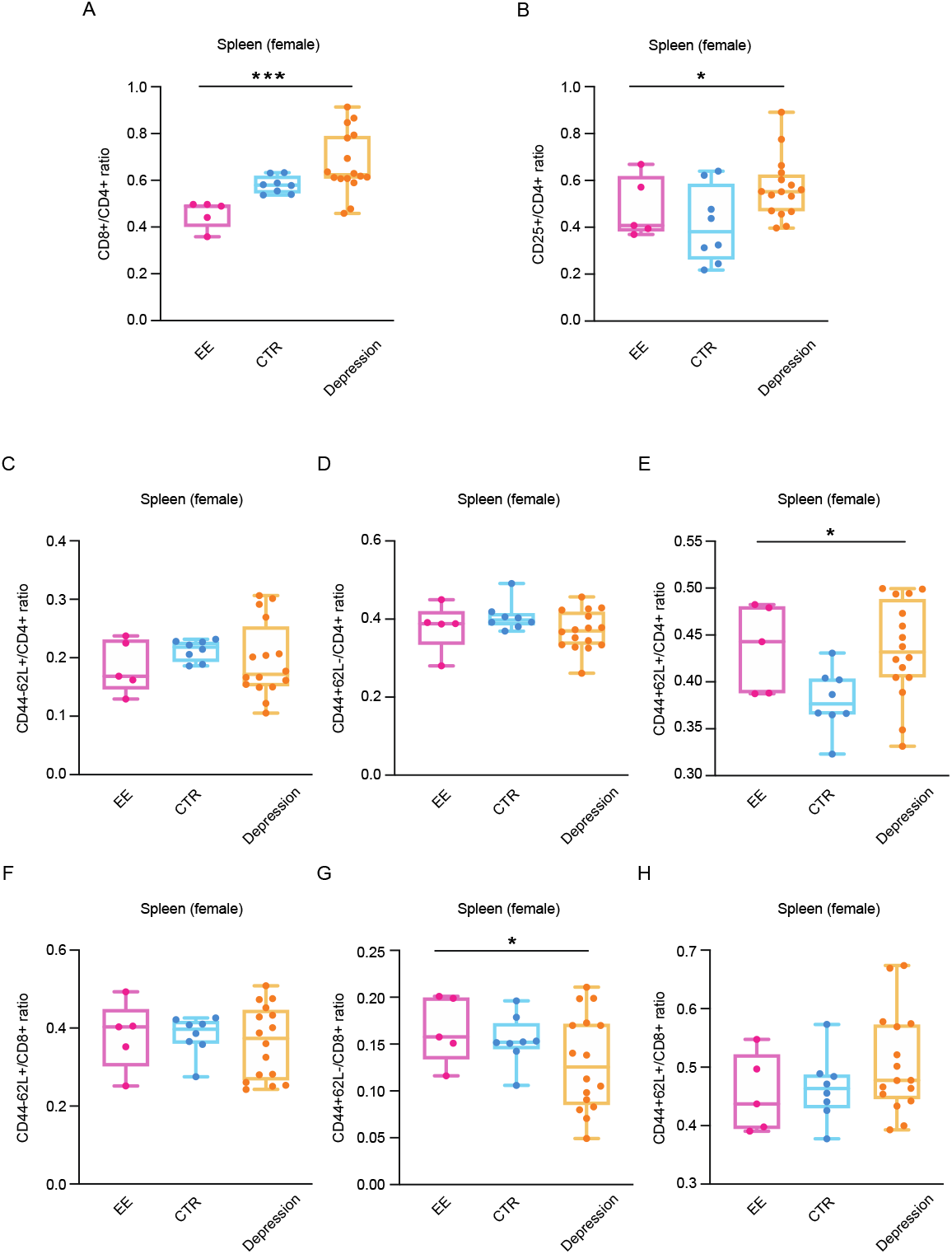
T cell populations in spleens of 10-12-month female mice. **A,** Statistics of CD8+/CD4+ ratio in the spleens of mice across three groups. **B,** Statistics of CD4+CD25+ (regulatory T cells)/CD4+ ratio in the spleens of mice across three groups. **C-E**, the ratio of naive T cells (CD44-CD62L+), effector memory T cells (CD44+ CD62L-) and central memory T cells (CD44+ CD62L+) of CD4+ T cells in the spleens of mice across three groups. **F-H,** the same as **C-E,** but data from CD8+ T cells. Ordinary one-way ANOVA test for more than two groups; * for p < 0.05; ** for p < 0.01; *** for p < 0.001; **** for p < 0.0001.

**Figure S6.**
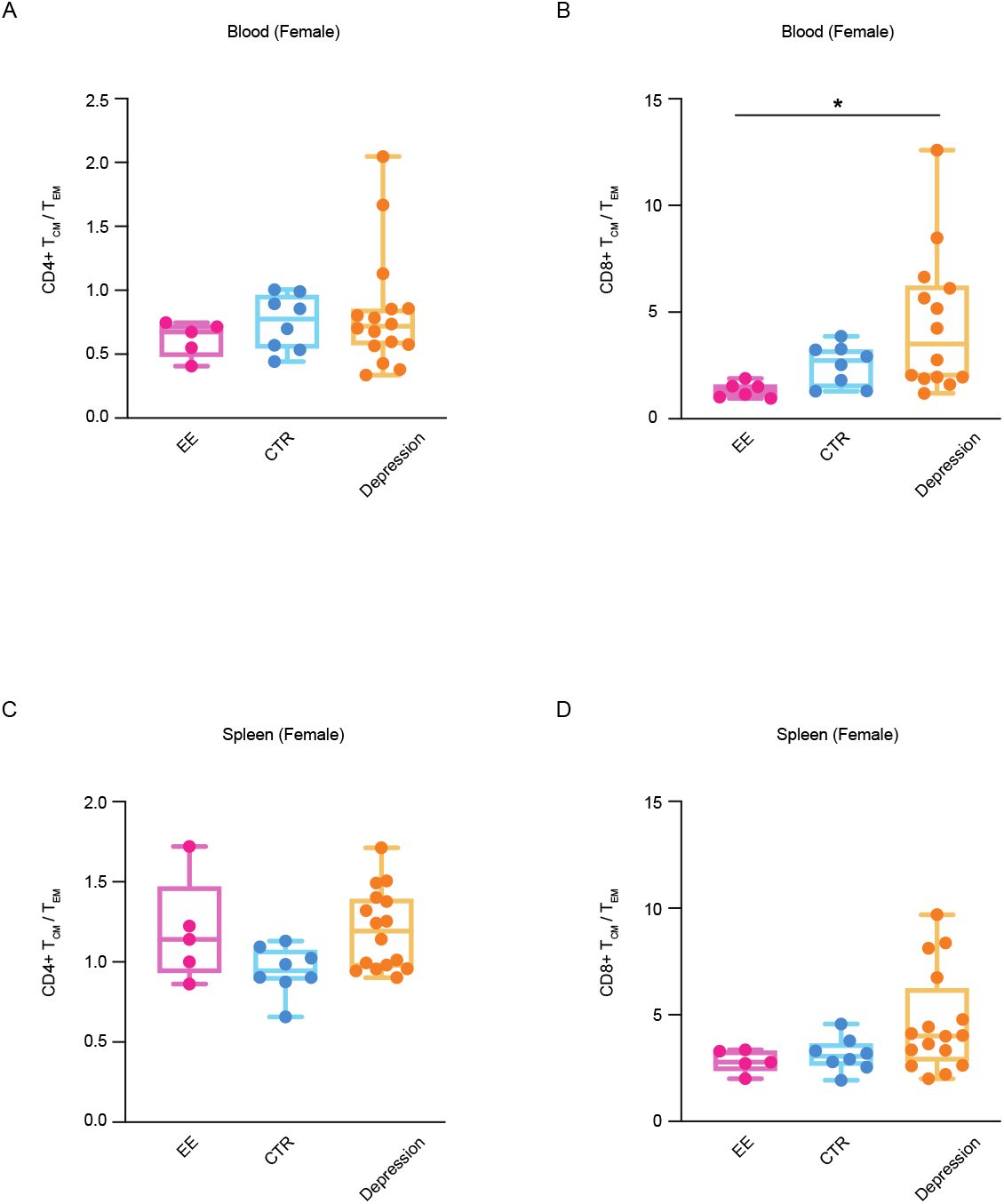
T_CM_-T_EM_ ratio of 10-12-month female mice. **A,** CD4+ T_CM_-to-T_EM_ ratio of peripheral blood. **B,** CD8+ T_CM_-to-T_EM_ ratio of peripheral blood. **C, D**, the same as **A, B** but statistics for spleens. Ordinary one-way ANOVA test for more than two groups; * for p < 0.05; ** for p < 0.01; *** for p < 0.001; **** for p < 0.0001.

## Notes

### Competing Interest Statement

The authors have declared no competing interest.

